# Does a strong reduction of colony workforce affect the foraging strategy of a social pollinator?

**DOI:** 10.1101/622910

**Authors:** Paolo Biella, Nicola Tommasi, Asma Akter, Lorenzo Guzzetti, Jan Klecka, Anna Sandionigi, Massimo Labra, Andrea Galimberti

## Abstract

The way pollinators gather resources may play a key role for buffering their population declines. Social pollinators like bumblebees could adjust their foraging after significant workforce reductions to keep provisions to the colony optimal, especially in terms of pollen quality, diversity, and quantity. To test what effects a workforce reduction causes on the foraging for pollen, colonies of the bumblebee *Bombus terrestris* were experimentally manipulated in field by removing half the number of workers. The pollen pellets of the workers were taxonomically identified with DNA metabarcoding, a ROC approach was used to filter out underrepresented OTUs, and video cameras and network analyses were employed to investigate foraging strategies and behaviour. The results suggested that the plant diversity in the pollen pellets was high but plant identity and pollen quantity traits were influenced mainly by plant phenology. During the experiment, although the treated colonies increased foraging effort in relation to control nests, only minor changes in the diet breadth and in the other node-level and network-level indices were observed after workforce removal. Therefore, a consistency in the bumblebees’ feeding strategies emerges despite the lowered workforce, which questions the ability of social pollinators to adjust their foraging in the field.

## Introduction

Social pollinators, such as bumblebees, are subjected to multiple stressors that ultimately cause population reductions. These declining trends are mostly due to climate change^1,2^ and several “pollinator-unfriendly” practices related to agriculture (i.e., a general intensification, the use of monocultures, the use of harmful agrochemicals^3,4^, and the use of synthetic fertilisers causing shifts in the vegetation^5^). Moreover, land use change^6^, the lack of flower diversity^7^ (e.g. overgrazing or frequent mowing^8^), the reduction of natural ecosystems nearby fields^9^, the spread of parasites and diseases^10^, and the overwhelming competition from domesticated bees^11,12^ also impact the dynamics of bumblebees and other pollinators’ populations.

Gathering sufficient and appropriate resources is a key nutritional aspect for stabilizing pollinator populations^13–16^. For pollinators whose development relies exclusively on plant pollen and nectar, the nutritional profile of the resources should eventually influence the way foraging choices are performed^17–20^. In other words, a bee should maximize the micro macronutrients of the resources it collects in order to provide a balanced and optimal diet to the developing brood^21^. However, it is not known how population declines could modify the way pollinators gather resources. According to the Optimal Foraging Theory, it can be expected that individuals are influenced not only by the reward’s nutrients but also by competitive interactions, so a higher forager density causes faster depletion of the resources and triggers a wider diet breadth in the foragers as consequence (i.e. density-dependent mechanisms ^22,23^). On the other hand, a significant loss of workforce in a social pollinator colony could lead to a sudden decrement of diversity and quantity of resources incoming to the nest. As a consequence to this, the colony should react in an adaptive and/or optimal way (i.e. gathering resources for enhancing and/or maximizing fitness)^24^. In other words, according to the Optimal Foraging Theory^25^, the colony could respond by augmenting the overall foraging effort to increase the amount of incoming resources, or the foragers could favour plants with high pollen production; alternatively, after workforce reduction, the foragers could enhance resource heterogeneity in order to assure the nutritional value given to the larvae by compensating for the resource types that had previously been brought into the nest by the foragers that went missing.

In the case of pollen collected by pollinators, studying insect-plant interactions is complicated by several methodological aspects. In addition to the direct observation of an insect’s behaviour^26^, the analysis of pollen on an insect’s body can reveal the interactions that happened during a pollinator’s trip and can also yield the rarest interactions that normally remain undetected during observational surveys^27,28^. Yet, morphology-based identification of pollen lacks a uniform discriminatory power and requires great taxonomical knowledge^29–31^. However, the potential benefits of pollen studies highlight the need to improve methods that are alternative to the morphological analyses. In this context, DNA-based approaches, such as DNA barcoding and DNA metabarcoding, represent reliable approaches^32,33^. In other words, by using integrative approaches (e.g. DNA metabarcoding applied to ecological questions), methodological issues can be overcome and the interactions and the resource usage by declining pollinators can be explored in more depth.

In this work, we tested the possible expectations about changes in foraging preferences due to colony workforce reduction by experimentally inducing a sudden decline in the colony size of commercial colonies of the bumblebee *Bombus terrestris* (Linnaeus, 1758) and by investigating consequent changes in foraging. We intended to recreate a situation of workforce loss due to natural or human-based environmental conditions (see ^34–36^). We explored the foraging behaviour before and after the manipulation with video recordings and also resource utilization by identifying the pollen with a DNA metabarcoding approach. We focused on bumblebees, because (a) they are among the most effective pollinators^37^, (b) they are social pollinators native to several regions of the world, while honeybees are present often due to domestication^38^, (c) their colonies need a high amount of resources which makes them dependent on the habitat they live in^2^, and (d) their natural populations are declining^1,10^. Our specific aims were to investigate the effect of an experimental reduction of the bumblebees’ workforce by focusing on responses (i) in the foraging strategies of individuals and in the associated bumblebee-plant networks, (ii) in the foraging rate per unit of time, and (iii) in the diversity of the collected plants and in plant’s traits of pollen production. This experiment has the potential of providing new insights into the ways social pollinators respond to environmental or anthropic events by interacting with plant resources within the context of pollination ecosystem services.

## Material and methods

### Study area, experimental set-up, and sample collections

The experiment was conducted in a meadow near Český Krumlov, 18 km southwest of České Budějovice (Czech Republic, 48°49’30.52’ N, 14°19’4.02" E), that belongs to a 62 ha natural area located at an altitude of 600 m a.s.l. and consists of forest, isolated trees, and shrubs, while a portion is covered by species rich calcareous grasslands managed by occasional extensive grazing. Around this zone, a mosaic of agricultural areas and urban settlements occurs. The study site is part of a publicly accessible area where sample collections are allowed (with the exception of species protected by law). The experiment and the collection of samples were carried out on sunny days without strong wind or rain, in summer 2016.

Four commercial colonies of the bumblebee *Bombus terrestris* were bought from a private company (Koppert s.r.o., Nove Zamky) and were placed in pairs at the study site at a distance from each other of about 500 m in order to capture possible minor changes in floristic composition. All colonies were marked and placed in the field under shade to prevent overheating. The number of used colonies lies within the range used in other studies about bumblebees foraging^18,39–41^. In each pair, a colony was used as a control, and it was not treated during the length of the experiment, while a second colony was used to apply a treatment of diminishing the worker population, which in practice consisted of manually removing 50% of the workers relative to the number of workers present in the period before removal in that colony. This removal threshold was inspired by studies reporting mortalities or worker losses up to 50% with respect to control colonies due to multiple stressors (see ^34–36^). For removing the workers, as we used nest boxes with a way-in and a way-out holes, the way-in was left open for an entire afternoon so that workers could return to the nest but none could leave it, and then the nest was completely closed during the following night. Early in the next morning, light anaesthetization with CO2 was applied to the nest for a very short time, workers were counted and half of the worker amount was removed from the nest.

Four days after placing the colonies in the field, the workers’ pollen pellets were collected from the corbiculae of the legs just before entering the nest and after light anaesthetization with CO2^42^ (the workers were afterwards released outside their nest to avoid immediate complications for the larvae related to workers being anesthetized^43^). The pollen of 18 bumblebee workers for each nest were surveyed before workforce halving (“before” phase, 6^th^−11^th^ July). In the period after removing the workers (“after” phase, 20^th^−23^rd^ July), pollen pellets of 18 workers for each colony were collected in the same way as the “before” period (17 workers for one of the nests). The number of samples collected was similar to other studies on DNA metabarcoding of pollen^44,45^. Pellets were collected with sterile tweezers and placed in Eppendorf tubes, marked with codes and stored in a freezer at −20 °C. The number of samples included in the analyses provided a plant diversity per nest that was estimated to be 83% and 78% of the asymptotic plant diversity of each treated nest as shown by the Chao2 estimator calculated by the *iNEXT* package of *R* with incidence data (±4 species in nest 1 and ±4.8 species in nest 2).

Local botanists provided an accurate check-list of the flowering plant species at the study area (i.e., 112 plant species, see Supplementary Information Table S1). Those species that were not available in public nucleotide databases (i.e., NCBI and BOLD) were sampled (i.e., 54 plant species, one or two young leafs each, stored at −20 °C) to create a complete DNA barcoding reference dataset. Reference ITS2 sequences for the remaining species were directly retrieved from GenBank NCBI prior to accurate validation of the accessions (i.e., availability of voucher details and complete overlapping with the DNA barcoding region sequenced in the bumblebees’ pollen pellets). Overall, the final reference dataset encompassed 1196 ITS2 sequences.

### DNA analyses and taxonomical assignments

Reference ITS2 DNA barcodes for the sampled plant species were obtained as described in ^46^ and deposited in EMBL GeneBank under the accessions reported in Supplementary Information Table S1.

For each bumblebee, one pollen pellet was grinded with a Tissue Lyser LT (Quiagen, Hilden, Germany) prior to freezing the sample in liquid nitrogen. The total DNA was extracted using the EuroGOLD Plant DNA mini kit (EuroClone, Pero, Italy), following the manufacturer’s instructions and using a final elution volume of 70 μl.

The identification of the plant diversity within pollen loads was performed through a HTS (High-throughput sequencing) DNA metabarcoding approach targeting the nuclear internal transcribed spacer 2 region (ITS2). This locus was successfully used for characterizing pollen-mixed samples in several recent studies^32,33,47^. DNA libraries for each sample were prepared following Illumina guidelines (16S Metagenomic Sequencing Library Preparation, Part #15044223 Rev. B) with modifications for ITS2 sequencing. The ITS2 region was amplified using primers S2F and S3R^48^ with the addition of the Illumina overhang adapter sequences, namely

S2F_Seq:

5’TCGTCGGCAGCGTCAGATGTGTATAAGAGACAGATGCGATACTTGGTGTGAAT 3’

S3R_Seq:

5’ GTCTCGTGGGCTCGGAGATGTGTATAAGAGACAGGACGCTTCTCCAGACTACAAT 3’.

Before amplification, DNA extracts were normalized by means of quantitative real-time PCR (qPCR) Ct values with the same amplification primer pairs and the same protocols described in ^49^. PCR reactions contained 12.5 μl of KAPA HiFi HotStart ReadyMix PCR Kit, 5 μl of each primer 1 μM (forward and reverse) and 2.5 μl DNA (maximum volume of DNA per sample with 5ng/μl DNA concentration). Samples were initially denatured at 94° C for 5 min, then amplified using 40 cycles at 94° C for 30 s, 56° C for 30 s, and 72° C for 45 s. A final extension (72°) of 10 min was performed at the end of the programme to ensure complete amplification. All PCR amplifications were prepared under an UV PCR cabinet to avoid contamination. The success of amplification was tested on a 1.5% agarose gel-electrophoresis. A 100 bp mass ladder (GeneDirex 100 bp DNA Ladder RTU, FroggaBio Inc., Toronto, ON, Canada) was used to confirm the successful normalization of the amplicon concentration within the samples.

Index PCR and library sequencing were performed through the Illumina MiSeq instrument using MiSeq Reagent Kit v3 (2 × 300-bp paired-end sequencing). The library preparation and the sequencing process were conducted at BMR Genomics (Padova, Italy). Raw Illumina reads were paired and pre-processed using *USEARCH* 8.0.1623 ^50^. Reads were filtered out if ambiguous bases were detected and lengths were outside the bounds of 250 bp. Moreover, an expected error of 1 was used as an indicator of read accuracy. OTUs (Operational Taxonomic Units) were obtained using -- *cluster_fast* algorithm from *VSEARCH.2* software (https://github.com/torognes/vsearch)^51^ with a 99% sequence identity. The cluster centroid was chosen as the representative sequence of the cluster. The taxonomic assignment of the representative sequences was carried out using the *BLAST* algorithm^52^ against the reference DNA barcoding dataset of the study area (see above), accepting only assignments with Max Identity and Query Coverage > 98%. OTUs representative sequences showing assignment values of Maximum Identity and Query Coverage < 98% with the use of this database were assigned using the GenBank NCBI database with the above-mentioned thresholds. Taxonomic assignment at a genus level were preferred instead of a species level if the queried OTUs resulted in a Max Identity and Query Coverage > 98% with several species of a given genus and co-occurring at the study site (in the case of the DNA barcoding reference dataset) or within the investigated geographic region, i.e South Czech Republic (in the case of NCBI queries).

### Selection of OTUs

Sorting false positives from data produced with DNA metabarcoding has been recently underlined^53^. In order to exclude false-positive OTUs from the dataset, the ROC (Receiver Operating Characteristic) framework was used to quantify a trade-off of acceptance or rejection of OTUs within the analyzed pollen samples. The ROC framework assesses the true positive rate and the true negative rate of a test^54^, based on the Youden index. This approach can improve the reliability of OTU assignments by establishing defensible thresholds for rejection or acceptance^55^. This is a well-accepted methodology for threshold detection, since it is used in several biological fields, including DNA- and environmental DNA-based studies^55,56^.

In the samples of this study, some OTUs were represented with only a very low number of reads. This would hint at the presence of false positives. Therefore, in order to find reliable thresholds, we followed the suggestions of ^55^ which employs ROC curves instead of arbitrarily cut-off values for excluding OTUs from the samples.

Specifically for each sample independently, a categorical variable “negative” was assigned to the OTUs with 0 number of DNA reads and “positive” was assigned to the OTU with reads >0. A GLM (Generalized linear regression) with an overdispersed Poisson distribution (quasipoisson) was performed independently on each sample in order to estimate the distribution of reads related to positives and negatives; the amount of reads per OTU was the response variable and "positive" or "negative" was the predictor variable. On the values estimated by the regressions, the *pROC* package^57^ in the R environment^58^ was used to estimate the per-sample cutting threshold and thus to identify which OTUs were false positives. Those OTUs with a number of reads below the estimated thresholds were excluded from the dataset and considered as false positives. The resulting dataset was used in the following analyses (Supplementary Information Table S2, Supplementary Dataset Table S3).

### Networks of foraging

For each nest and at each experimental phase (time “before” and “after” removal of workers), matrices representing the interactions of bumblebees and plants were analysed to investigate changes in the foraging strategies by means of networks analyses (data in Supplementary Dataset Table S3). Both binary and quantitative matrices were used, because different aspects are accounted for. The binary ones are useful for studying network structures where all links are equal (as they are based on the presence and absence of interactions), while in the quantitative ones, the links have different weights according to the intensity of each interaction (e.g. interaction frequency, transferred biomass, etc).

Firstly, we tested several node-level indices (where a “node” is either a foraging bumblebee or a plant). Specialization was investigated using: (a) the “degree”, that is the number of plant species found in a pollen pellet; (b) RR, the “resource range”, that estimates the fraction of used resources to the total available^59^ and is computed here as 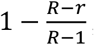, where *R* is the available resources (= plants) and is the used plants; (c) PG, the “proportional generality”, is the quantitative diversity of consumers in relation to the potential resources; it is computed as the ratio between the power of the quantitative Shannon diversity for consumers and that for the abundances of resources *q*:*e^H_p_^*/*e^H_q_^*; (d) PDI, the “Paired Difference Index”, is the quantitative counterpart of RR and it compares the strongest quantitative interaction with all remaining interactions^60^; it characterizes the decay of performance as drift from the optimal resource; it is calculated here as 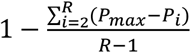, where *P_ma_* is the maximum of all quantitative interactions, *P_i_* is the quantity of interaction with the plant, and *R* is the number of available resources (=plants); (e) *d*’ index, which is a measure of specialization based on niche overlap among nodes^61^ and is calculated as 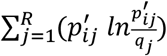, where *R* is the number of resources, *p′_ij_* is a species’ *i* interaction with partner *j* as proportion of the sum of interactions of *i,q_j_* is the sum of interactions of partner *j* divided by the total of all interactions. In addition, we studied the importance of plant species in the foraging network with the (f) “closeness centrality”, which indicates how a plant is near the core of the interactions based on the path lengths of the network; 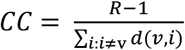, where *R* is the available plants and *d*(*v,i*) is the geodesic distance between plant and *i*^62^. Indexes (a), (b), and (f) are calculated from the binary interaction matrices (presence or absence of a plant in a sample), while indexes (c), (d), and (e) are based on the quantitative interaction matrix including the number of DNA reads of a certain plant species in a pollen pellet. Using DNA reads as a proxy of a quantitative amount of pollen was decently supported in Bell et al. (2018) and was already applied to networks in Pornon et al. (2017); these indexes include normalizations by matrix total. For testing changes in these indexes, each one was analysed with generalized linear mixed-effect models with library *lme4* ^63^ in the R environment with a given index as response variable, treatment as a predictor variable (“before”, “after” worker removal), and nest identity as the random intercept. Poisson distribution or Gamma distribution with the log link function were used, depending on the response variable.

Secondly, to test whether the entire bumblebee-plant networks changed after the treatment, the interaction matrices included either binary interaction matrices or the count data of the DNA reads, such as in, standardized^28^ by the total of the matrix. For each nest, the network structure was studied by focusing on several aspects of networks. Firstly, the proportion of realized interactions was studied with (a) Link density LD^64^, which is a quantitative measure of the proportion of realized interactions weighted by interaction diversity and is computed as 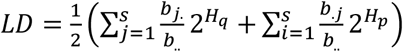, where is the number of species in the networks, is the total sum of the matrix, is the sum of the interactions of bumblebees and is the sum of the interactions of plant *i*, *H_q_* is calculated as 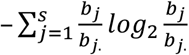 with as an interaction (and similarly for plants and plant species *i*); (b) Connectance C ^64^, which is the proportion of realized links in the network and is calculated as 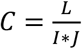, *L* is the number of interactions, *I*, and *J* and is the number of plant and animal species, respectively, and can vary from 0 to a maximum of 1. In addition, how resources are distributed among nodes was investigated with the nestedness index, so that in a nested network the generalist pool interacts with both specialists and generalists. It was calculated as (c) Nestedness based on Overlap and Decreasing Fill (NODF) and (d) the weighted counterpart WNODF^65^, is based on decreasing fill and on paired overlap on the matrix. Between pairs of columns and pairs of rows, it detects the degree of nestedness by comparing the marginal totals and the proportion of filled matrix cells located at the same position. Thus, for a matrix with plants and bumblebees, 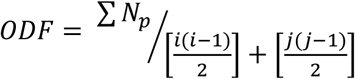. It ranges from 0 to 100 (fully nested). Moreover, the tendency of the network to divide into compartments, with implications for resource accessibility and competition, was calculated as (e) Modularity, and (f) the quantitative counterpart, computed by the algoritm *DIRTLPAwb*+ ^66^; *Q* is computed as 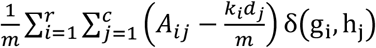, where *A_ij_* is the interaction matrix of rows and columns, is the number of links, is the node degree for a plant with label *h*, and *d* is the node degree for a bumblebee with label *g*, while the Kronecker function δ(g_i_, h_j_) is 1 if nodes and belong to same module or 0 otherwise. and range from 0 to its maximum of 1. Network-level specialization was also investigated. A niche-overlap measure of specialization of network-level interactions was studied with (g) Interaction Diversity 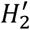. It is computed as 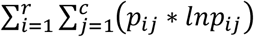, with and referring to rows and columns of the interaction matrix between a plant species and pollinator species, and is the proportion of the number of interactions in relation to the respective row total. Its possible maximum and minimum are obtained from the distribution of interaction totals of the matrix and used to normalize the index to vary between 0 and 1 (perfect specialisation)^61^. Specialization was also studied with (f) Generality and (g) Vulnerability indexes, that are the mean effective numbers of partners, that is of plants for bumblebees (Generality G) and of bumblebees for plants (Vulnerability V), weighted by the marginal totals; they are calculated as 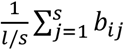, where a node is interacting with a node *j*, and *b_ij_* is the sum of quantitative interactions between *i* and *j*, and the total number of links in the network is and that of nodes is *s*^64^.

Changes in these indices of network structure were tested by means of random permutations of the data, which test whether the difference between the observed networks is significant with respect to random expectations. To reach this goal, the interactions (matrix cells) of both networks were swapped randomly between the two networks (“before”, “after”), following^67,68^, for 10000 times for each of the two networks. After each swap, the value of the difference was recalculated. The statistical significance was obtained by comparing the observed difference to the distribution of differences from the random permutations.

The node and the network indices were calculated with the packages *bipartite*^69^ and *vegan*^70^ in R.

### Foraging rate

All colonies (treated and control) were recorded with video cameras (Canon Legria HFR56) for a sample of three hours during a day during the experimental phases before removing workers and after removing workers. The camera was placed near the entrance of the nests, so that the number of leaving and of returning workers could be counted.

For testing changes in the foraging rate, the number of workers leaving during each 20 minute interval time was used as response variable in generalized linear mixed-effect models with library *lme4* in the R environment. The experimental phase was the predictor variable (i.e., the period “before” and “after” worker removal), and the nest identity was the random intercept. Poisson distribution with the log link function was used. An offset with the number of workers leaving the control nests per 20 minute time was included in the analyses, in order to account for variations of foraging rate independent from the treatment.

### Plant diversity in the pollen pellets

To investigate changes in plant species composition in the pollen samples during the experimental time, a PER-MANOVA (Permutational Multivariate Analysis of Variance Using Distance Matrices) was used with the function *adonis* in the *vegan* package in R. Samples-per-plants matrices with presence / absence of a given plant in a given sample was considered the response variable, while the experimental phase (treatment of “before” and of “after” removal of workers) and nest were predictor variables. Treated and control colonies were analysed separately.

### Traits of pollen production

Values of the plant trait of pollen production (“pollen quantity”) were assigned to both used and unused plants flowering at the study area during the experimental time (Supplementary Information Table S1 and Table S2). These values were extracted from ^71^, which ranks plants from low (“P0”) to high (“P5”) levels of pollen quantity in several European plant species. Specifically, this ranking is based on the amount of pollen produced by the plant species, it is coherent with other rank-based pollen databases (see ^72^) and it also is incorporated in the plant trait databases provided by the *TR8* package for R^73^. We deem that pollen production is a suitable plant trait for the comparative purpose of testing changes in food preference before-after a treatment in bees.

The probability of collecting pollen of a species was analyzed using logistic regression (generalized linear mixed-effect models) with presence/absence of a plant in a sample for a given colony as a response variable, treatment in interaction with (numerical) pollen quantity as predictors and nest identity as a random intercept, with binomial distribution and logit as link function, with the *lme4* package for R. Confidence intervals were estimated with 1000 bootstrapping using the function *bootMer*. Control and treated nests were analyzed separately.

## Results

### Sequencing, filtering, and taxonomic assignment of pollen loads

Illumina sequencing of pollen samples yielded 18,473,760 raw reads. After pair-ending and quality filtering, 5,600,000 reads were included in the dataset, and they were clustered in 167 OTUs, 51 of which showed high similarity with fungi accessions and were excluded from the dataset. The remaining OTUs were assigned to 44 plant taxa and specifically 90 OTUs (53.9%) to the species level and 26 OTUs (15.5%) to the genus level. The ROC filtering excluded 25 additional OTUs (at least 10 plant species) and a total of 72,361 reads were removed from the dataset with a mean of 36 reads per sample (with across sample st. dev. of 101 reads and a range of 0 – 1214 reads) which corresponds to an average of cutting thresholds across samples of 2.28% of the reads. Therefore, the filtered list of plant species encompassed 34 taxa (91.2% with species identity) with a mean of 2.25 taxa per sample, st. dev. = 1.54, min. 1 and max. 10 (Supplementary Information Table S2 and Supplementary Dataset table S3), corresponding to 3,392,081 total reads.

Ten taxa in the post-ROC dataset were not initially included in the floral checklist. Monofloral pollen pellets were 37% (53 samples), while 63% (90 samples) were polyfloral (44 pollen samples of two plant taxa, 46 samples of more than two taxa).

### Pollen plant diversity, node- and network-level responses to the treatment

Taxa composition of the pollen samples changed over the study period, both in the control and in the treated nests (Supplementary Information Figure S1). In both the treated and control colonies, the experimental phase (before/after workforce reduction) predicted the plant identity of the pollen samples better than the nest identity, although both variables were significant (Table 1).

**Table 1.** Results from the PER-MANOVA statistics testing the effect of worker removal and nest identity in the presence/absence of plant species in the pollen pellet samples. Significant results are highlighted in bold.

|  | Variable | Df | R<sup>2</sup> | P |
| --- | --- | --- | --- | --- |
| Treated<br>nests | Experimental phase | 1 | 0.108 | **0.001** |
|  | Nest Id. | 1 | 0.068 | **0.001** |
|  | Residuals | 68 | 0.825 |  |
|  | Total | 70 | 1 |  |
| Control<br>nests | Experimental phase | 1 | 0.111 | **0.001** |
|  | Nest Id. | 1 | 0.061 | **0.003** |
|  | Residuals | 69 | 0.828 |  |
|  | Total | 71 | 1 |  |

The node level network analyses in the phase before removal revealed that Degree and PG were low but the plant’s Closeness Centrality was high and PDI and RR were both low, while d’ spanned over a wide range of the specialization-generalization gradient (Fig. 1). Changes after treatment were not significant, except for the quantitative PG index which changed significantly only in the control nests (Table 2 and Fig. 1).

**Table 2.** Individual level foraging indices tested for significant changes after halving the colony workforce, by generalized linear mixed-effects models. Statistical significance is highlighted in bold.

|  | Type | Treated nests | Control nests |
| --- | --- | --- | --- |
| (a) Degree | Binary | $\chi^2 = 0.11$, df=1, p = 0.74 | $\chi^2 = 0.53$, df=1, p = 0.467 |
| (b) RR,<br>Resource range | Binary | $\chi^2 = 0.897$, df=1, p = 0.344 | $\chi^2 = 0.157$, df=1, p = 0.692 |
| (c) PG,<br>Proportional<br>generality | Quantitative | $\chi^2 = 1.41$, df=1, p = 0.235 | $\chi^2 = 5.113$, df=1, p = **0.024** |
| (d) PDI, Paired<br>Difference<br>Index | Quantitative | $\chi^2 = 0.322$, df=1, p = 0.571 | $\chi^2 = 0.151$, df=1, p = 0.698 |
| (e) d',<br>Complementary<br>specialization | Quantitative | $\chi^2 = 0.2$, df=1, p = 0.655 | $\chi^2 = 0.2$, df=1, p = 0.648 |
| (f) Plant's<br>Closeness<br>Centrality | Binary | $\chi^2 = 1.28$, df=1, p = 0.258 | $\chi^2 = 2.092$, df=1, p = 0.148 |

**Figure 1.**
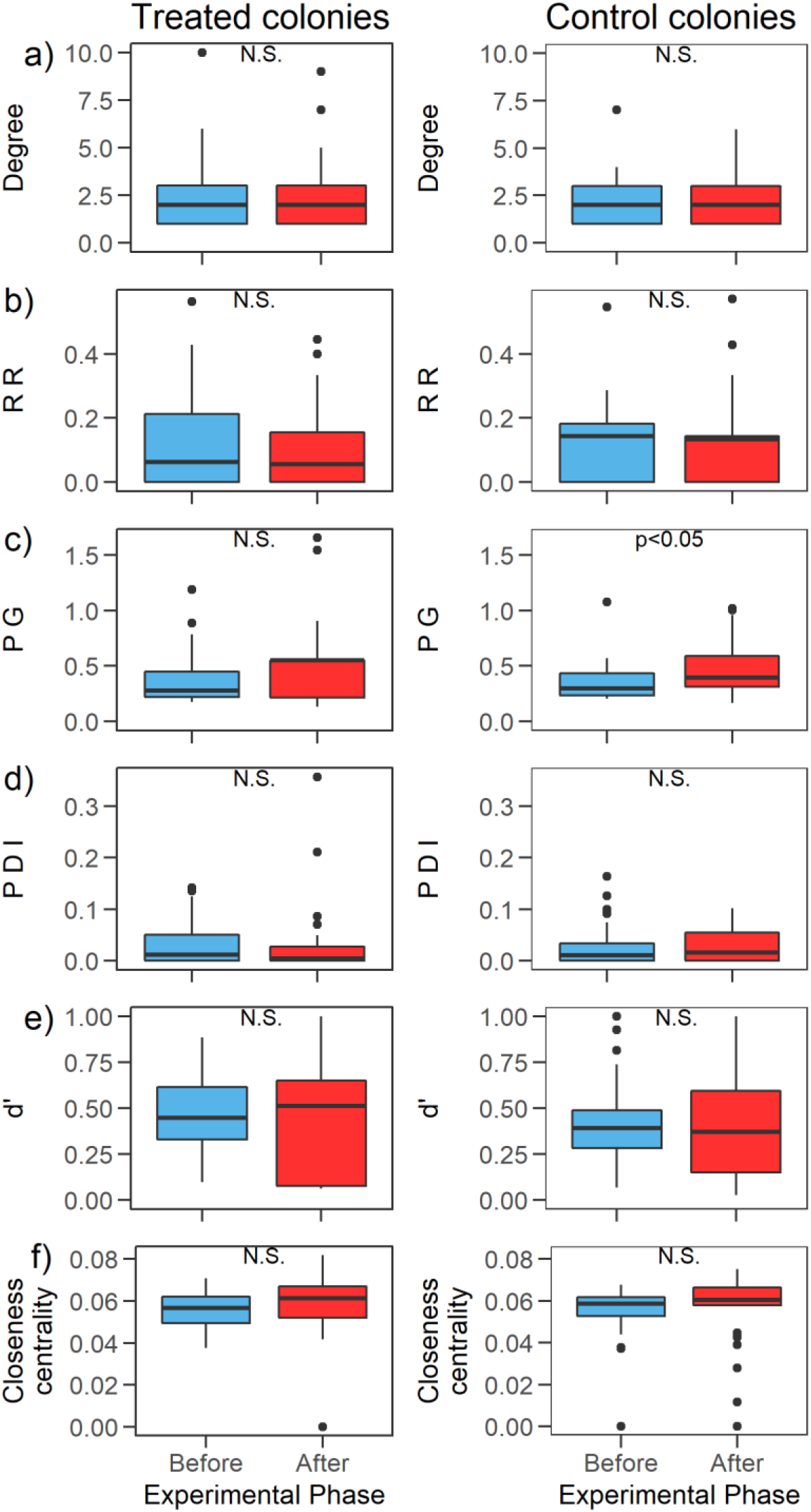
Node-level network indices of foraging and their change during the experiment: (a) Degree, (b) RR: Resource Range, (c) PG: Proportional Generality, (d) PDI: Paired Difference Index, (e) d’: Complementary specialization, (f) Closeness Centrality for plants (see methods). “N.S” signifies not statistically significant and the statistical tests are GLMMs (see methods).

The binary indexes of the network-level analyses didn’t change significantly after treatment in the treated colonies. On the other hand, only two of the quantitative indexes (i.e., the Link Density and Vulnerability of plants) changed significantly over the study period (Table 3 and Fig. 2).

**Table 3.** Network indices tested for change during the experimental phases (before and after the worker removal) by 10000 random permutational swaps of interactions between the networks before and after the treatment. Statistical significance is highlighted in bold. N1 and N2 indicate the treated nest’s identity.

|  | Type |  | Before | After | P | P control |
| --- | --- | --- | --- | --- | --- | --- |
| Link Density | Quantitative | N<sub>1</sub>= | 3 | 1.87 | **0.006** | 0.335 |
|  |  | N<sub>2</sub>= | 2.94 | 5.57 | **0.043** | 0.224 |
| Connectance | Binary | N<sub>1</sub>= | 0.17 | 0.15 | 0.221 | 0.05 |
|  |  | N<sub>2</sub>= | 0.25 | 0.25 | 1 | 0.677 |
| NODF | Binary | N<sub>1</sub>= | 17.21 | 11.55 | 0.054 | 0.482 |
|  |  | N<sub>2</sub>= | 7.68 | 7.96 | 0.926 | 0.625 |
| Weighted NODF | Quantitative | N<sub>1</sub>= | 12.77 | 12.45 | 0.925 | 0.881 |
|  |  | N<sub>2</sub>= | 8.33 | 5.18 | 0.613 | 0.96 |
| Modularity | Binary | N<sub>1</sub>= | 0.42 | 0.5 | 0.09 | 0.581 |
|  |  | N<sub>2</sub>= | 0.48 | 0.47 | 0.829 | 0.634 |
| Weighted Modularity | Quantitative | N<sub>1</sub>= | 0.65 | 0.75 | 0.151 | 0.248 |
|  |  | N<sub>2</sub>= | 0.68 | 0.26 | **0.015** | **0.046** |
| H2' | Quantitative | N<sub>1</sub>= | 0.85 | 0.92 | 0.365 | 0.362 |
|  |  | N<sub>2</sub>= | 0.87 | 0.83 | 0.768 | 0.31 |
| Generality | Quantitative | N<sub>1</sub>= | 1.67 | 1.47 | 0.388 | 0.089 |
|  |  | N<sub>2</sub>= | 1.34 | 1.17 | 0.381 | 0.969 |
| Vulnerability | Quantitative | N<sub>1</sub>= | 4.34 | 2.27 | **0.004** | 0.459 |
|  |  | N<sub>2</sub>= | 4.55 | 9.97 | **0.032** | 0.199 |

**Figure 2.**
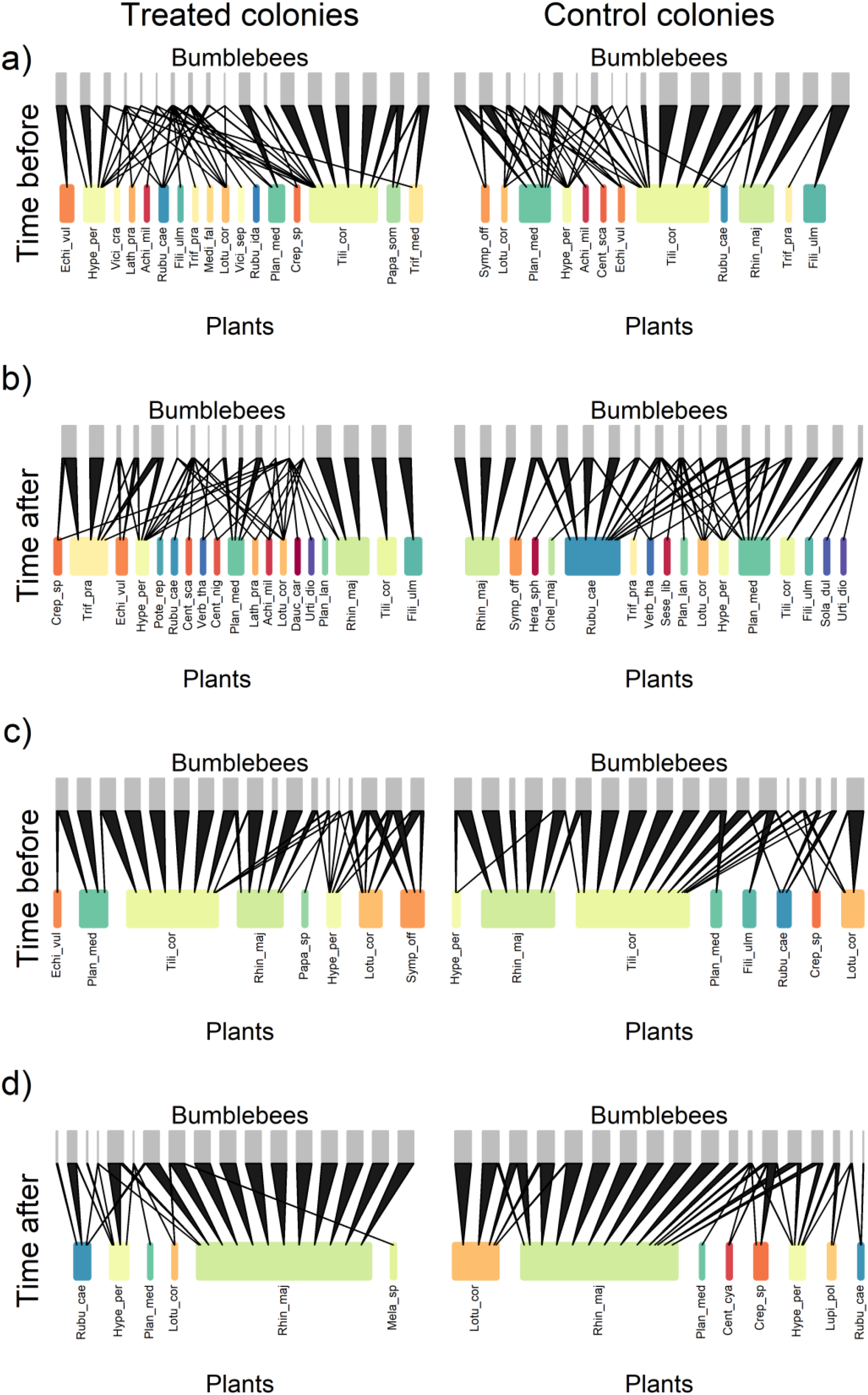
Bumblebee-plant networks during the experimental phases of before (plot’s panels “a” and “c”, first and second nest respectively) and after (plot’s panels “b” and “d”, first and second nest respectively) the workforce removal. For each plant species a colour is given and the plant’s full name is provided in Table S2.

### Foraging rate

After removing the workforce, the proportion of workers leaving the treated nests relative to the control nests’ foraging rate increased (Fig. 3, the trend without the proportion to control’s foraging is in Supplementary Information Figure S2). Specifically, the treatment was a significant predictor of the number of workers leaving in the GLMM with an offset of the control’s leavings (β_after_ – β_before_ = 0.40, likelihood ratio test χ^2^=14.945, df=1, p < 0.001).

**Figure 3.**
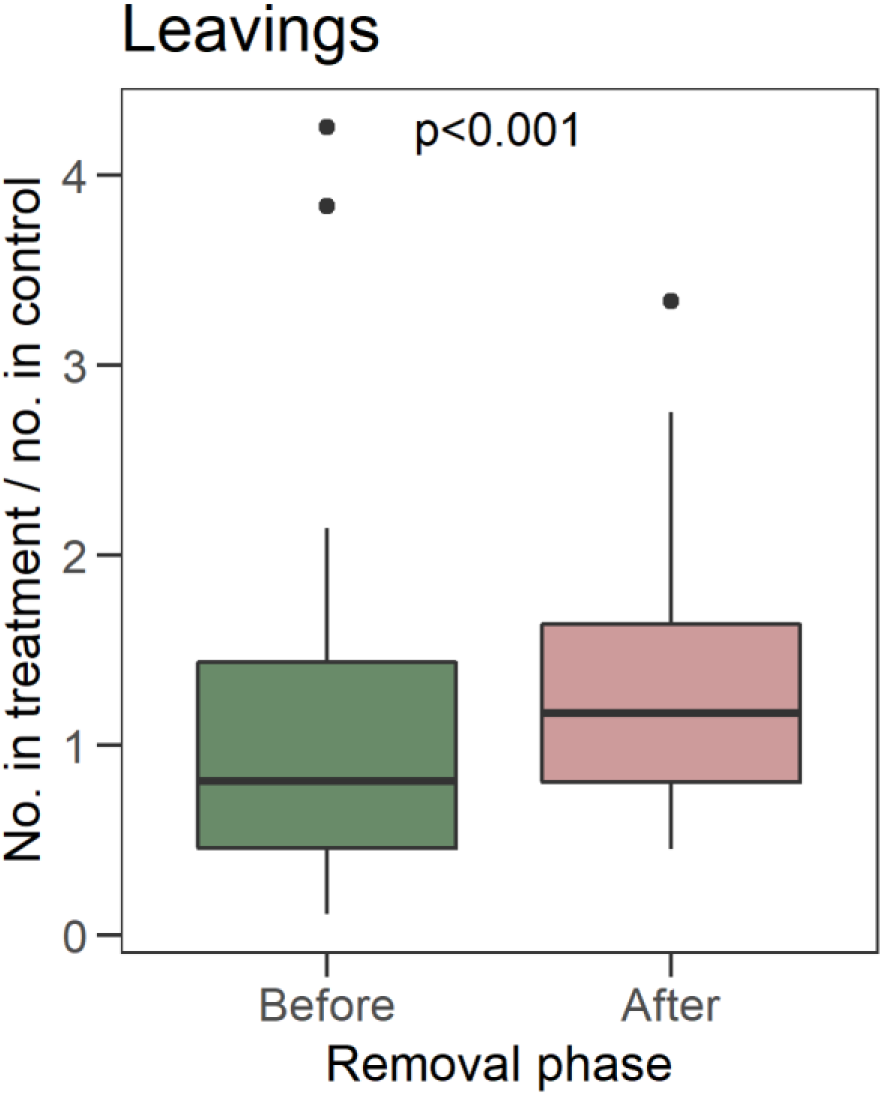
Number of workers leaving their nests per time unit (20 minutes long) proportionally to the control’s leaving during the same time units. Significance is tested with a GLMM (see methods).

### Pollen quantity

The probability of collecting plants of high pollen quantity decreased during the phase after the workforce removal with respect to the phase before, in both control and treated colonies (Fig. 4). The interaction of treatment and pollen quantity well predicted the collection probability (β_after_ – β_before_ = −0.55 and likelihood ratio test χ^2^=19.351, df=1, p < 0.001 in the treated colonies; β_after_ – β_before_ = −0.35, χ^2^=6.67, df=1, p < 0.01in the control colonies).

**Figure 4.**
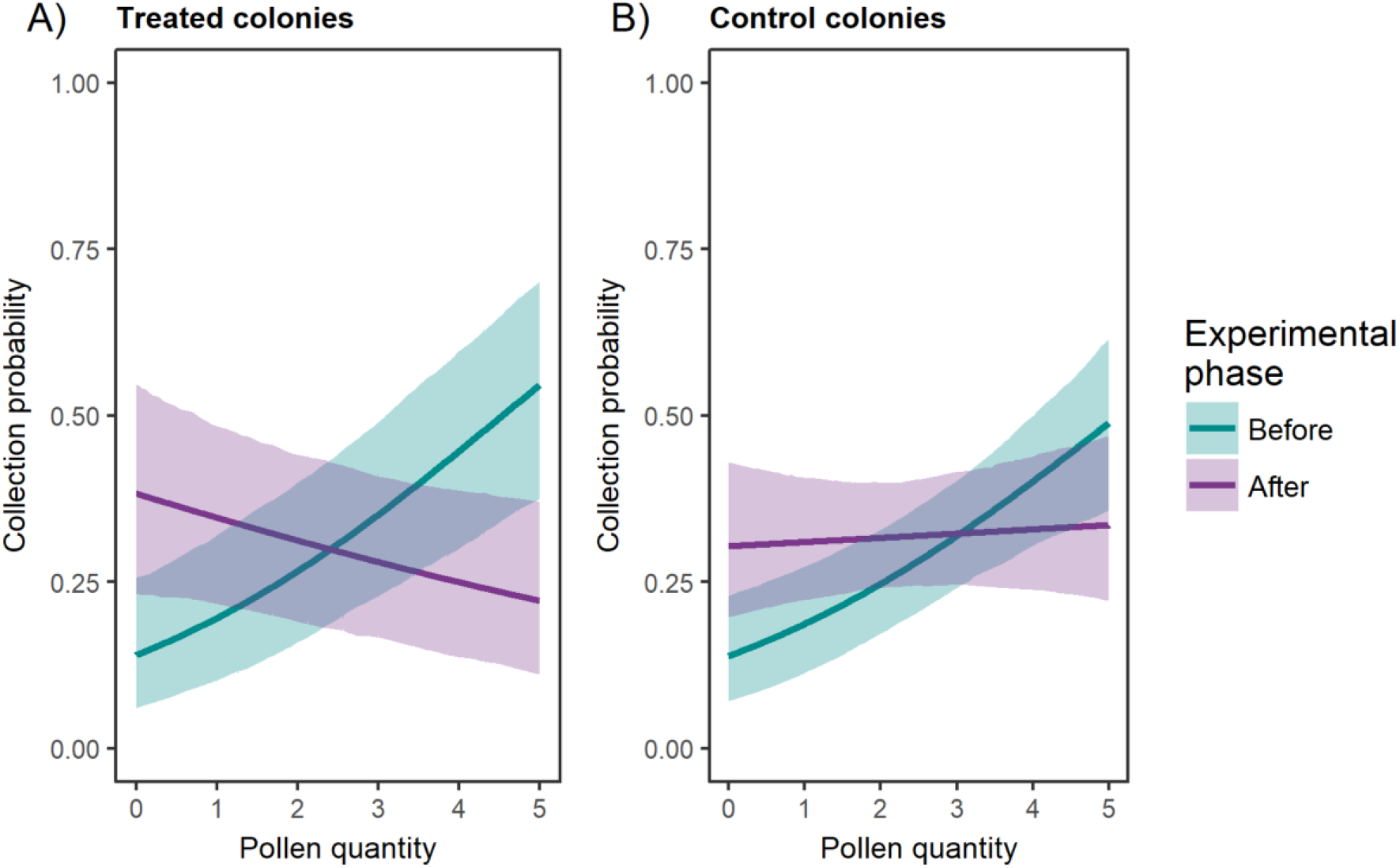
Pollen quantity of the foraged plants during the experimental phases in the treated (panel “A”) and control colonies (panel “B”). The plot shows the estimated probabilities (lines) and the 95% confidence intervals (polygons).

### Discussion

Previous studies on the foraging activity of bumblebees mainly focused on altering a diet and investigating adjustments in foraging in laboratory conditions^22,39,40^. A novel aspect of our study is that we investigated how reductions in colony size would affect the resource utilization and the foraging behaviour of these key pollinators when free to forage in the field. To our knowledge, only two studies have previously investigated the effect of experimentally removing the bumblebees’ workforce exclusively on colony fitness ^74^ and on the feeding of larvae^75^. Nevertheless, in our study we have focused on aspects related to foraging in the field and to the resource utilization by bumblebees when collecting plant pollen.

We acknowledge that our experimental design could have included more replicates (e.g., a higher number of colonies) and more samples of pollen and that these could strengthen the results. Even though our sample size and number of replicates was similar to other studies (see methods) and even though accumulation curves revealed that an acceptable level of plant diversity was yielded from the pollen samples, we encourage researchers to employ experimental designs that are replicated more. In our experimental design, we chose to of the pollen samples recovered from the bumblebees’ foragers were polyfloral, and this is consistent with the literature^77^. These are considered particularly beneficial for larvae alimentation in polylectic bees as the pollen is the main source of nutrients and secondary metabolites, such as vitamins and antioxidants^79^. Therefore, stocking polyfloral pollen pellets in the nest is considered to be an adaptive advantage for overcoming the among-plants variability of pollen quality^30^ and the risk of unbalanced and un-nutritious diets^79^, and also for strengthening the nutraceutical value of the diet^80^.

Plant choice was influenced by the pollen-production traits of the plants and pollen quantity played a role in resource utilization by the bumblebees as they preferred pollen from plants that were high-ranked in the pollen-production database we have used (Fig. 4). Nevertheless, in the experimental phase after workforce removal, less rewarding plants prevailed in the pollen samples, but this occurred both in the control and in the treated nests. We interpret this change as occurring due to factors other than the experimental manipulation of the workforce and, particularly, the slight phenological changes in the plant assemblage at the study site could have an effect on plant choices by the foragers.

That the vegetation phenological changes played a role is also supported by the fact that the workers from all nests, both treated and control ones, collected a diversity of plants that was different between the “before-removal” phase and the “after-removal” phase (Table 1, Supplementary Information Figure S1). Thus, it is possible that several plants shifted the status of the anthers’ maturation while still blooming during the time of the experiment, as it is common in plants^81^.

Despite the effect of these subtle phenological changes on the plants collected, the foraging strategies of individual bumblebees changed only slightly in the treated nests. In other words, the nodes-level network indices revealed small and non-significant changes in use a balanced sample size in order to rule out confounding effects due to varying sample sizes resulting from sampling more the colonies with higher foraging rates. Furthermore, we deemed that adding other study areas would be not practical due to the high heterogeneity of the landscape surrounding the location of the study and this could have added a confounding habitat effect to the outcomes of the experimental manipulation. In our study, the identification of pollen using DNA metabarcoding was reliable, because the plant list found in the pellets of our study matches other central European surveys^76,77^. Overall, the list of 34 plants found in the pollen samples retrieved from the individual foragers over the short time of our study highlights how polylectic bees normally rely on a wide set of flowering species^30,78^. Furthermore, it is possible that collecting more samples will yield a longer plant species list. Not only was the total plant diversity large, but our results also show that single foragers were indeed polylectic, as more than 60% specialism/generalism, in the number of gathered plants, in the proportion of the available resources actually collected, and in the centrality in the plants (i.e., the importance based on the position in the network) (Table 2, Fig. 1). Therefore, although the bumblebees changed the set of visited plants and their pollen-production traits, it seems that it did not change the way foragers utilized resources. This result is particularly surprising, because it contradicts the expectations based on density-dependent foraging that should have taken place due to an altered intra-specific competition after the manipulation. That is, a higher abundance of foraging bumblebees (as when workforce is high) can force foragers to use more plants and this implies a higher generalisation in resource use^22^. Conversely, our study did not find higher specialisation when foragers were few (i.e. during the “after” removal period in the treated nests). Our findings are supported by other choice experiments that showed a constancy in plant usage at higher bumblebee densities^41^ as well as that higher forager density did not change bumblebees’ foraging behavioural traits^39^. Therefore, whether or not pollinators forage according to density dependent mechanisms deserves further study in order to clarify how resources are collected in relation to forager density, at least under field conditions.

Furthermore, we expected to detect other compensating behaviours in the treated colonies, such as an increased foraging effort in the colonies that were subjected to the workforce removal. Actually, we have recorded that the foraging rate increased after workforce removal in the treated colonies, relative to control nests (Fig. 3), which suggests an increase in the foraging effort of the colonies. Increased foraging rates were recorded also in honeybee colonies after reductions in the amount of stored pollen^42,82^, which suggests a link between the foraging rate and the amount of pollen stored in the nest. Furthermore, the higher foraging rate we have found could either indicate that foragers made more foraging bouts per time unit and thus used more energy in travelling, or the alternative hypothesis of an increase in the number of foraging workers relative to colony size. Furthermore, the increased foraging rate could result in storing a higher amount of pollen in the nest or in storing an overall wider plant diversity, to compensate for the missing workers. A limitation of our study is that the pollen stored in the nests was not evaluated in spite of its potential of revealing deeper colony-level responses, because we did not want to cause disturbance to the treated or control colonies other than by removing workforce. Furthermore, we expected that bumblebees will collect more heterogeneous resources after workforce reduction and that this will impact the feeding networks of the bumblebees and the plants they collected pollen from. Conversely, our results suggest that the diet breadth did not expand and the bumblebee-plant network was not impacted by the workforce manipulation, as suggested by the indices of binary networks (based on presence/absence of interactions), and by the minor changes found in quantitative network indices related to the quantity of the resources used by foragers (Table 3, Fig. 2). This constancy in the foraging networks after workforce reduction can be explained by some aspects of bumblebees’ biology. In contrast to honeybees, the bumblebees are primitively eusocial which implies that colonies’ performance tends to rely more on individual choices of single foragers than on social information^40,83^ (the latter being the case of honeybees). This results from workers of *Bombus terrestris* having almost no contact with the larvae during their development^84^. Thus, foragers of our treatments might not have acquired any information on the development of the larvae and this could have prevented the expected foraging adjustment to take place. The lack of direct feedback between larvae and forager could uncouple the foraging choices and the colony’s growth rate, as it was clearly shown that removing workforce results in having less progeny and of smaller size^74^.

### Conclusions

By using DNA metabarcoding of pollen samples to overcome limitations of the morphological identification, this study investigated the effect of workforce decreases on the bumblebee foraging dynamics, on the chosen plant’s pollen-production traits and on the foraging rate, using an experimental manipulation in the field.

After applying a reduction of pollinator’s workforce, the bumblebees’ foraging strategies and the heterogeneity of collected resources were mostly constant, except for the increase in the colony’s foraging rate. If our results of a limited adaptation of foraging were confirmed by further studies and more replicated field-experiments, then these pollinators would have a limited ability to adapt to a decreased colony size as those that occur after multiple stressors (e.g., pesticide exposure, parasites, and diseases^85^).

## Acknowledgements

The authors thank K. Horvath for the linguistic revision, Antonella Bruno for her assistance in DNA libraries preparation and the botanists Alena Vitová, Petr Blažek, and Zuzana Chlumská. This study was supported by the Czech Science Foundation (projects GP14-10035P and GJ17-24795Y) and by University of South Bohemia (grant GA JU 152/2016/P). This survey was also founded by the ‘Ministero dell’Istruzione, dell’Università e della Ricerca’ (MIUR) within the project: ‘Sistemi Alimentari e Sviluppo Sostenibile – tra ricerca e processi internazionali e africani’; CUP: H42F16002450001. The funders had no role in conducting the research and/or during the preparation of the article.

## Author contribution statement

Conceived and designed the experiments: PB, JK. Performed the experiments: PB, AA, JK. Analysed the data: PB, NT, AS, AG. Drafted the manuscript and figures: PB, NT, AG, and ML. All authors contributed to the revision of the final manuscript.

## Data availability statement

All relevant data are within the paper and its Supporting Information files. The raw sequences obtained in this study were submitted to the European Nucleotide Archive (https://www.ebi.ac.uk/metagenomics). Study accession number is: PRJEB27433. Accession numbers of DNA barcoding sequences are available in GenBank NCBI system under the accession numbers LS973890-LS973943.

## Ethics statement

No permits were required for this project.

